# Effects of Spatial and Signal-Imposed Noises on Motor Unit Decomposition

**DOI:** 10.1101/2025.09.10.675422

**Authors:** Mansour Taleshi, Dennis Yeung, Francesco Negro, Ivan Vujaklija

## Abstract

High-density surface electromyography (HD-sEMG) decomposition offers insights into the neural drive through observation of individual motor units (MUs). However, ensuring that this method remains reliable under real-world signal degradations is crucial for its broader application. Therefore, we investigated the impact of three commonly modeled signal degradations on convolutive blind-source-separation (BSS) MU decomposition. 192 HD-sEMG channels were recorded from the forearm muscles of thirteen healthy participants during six wrist movements. Three broad categories of perturbation were introduced, including additive white Gaussian noise (WGN), channel loss, and electrode shift. These perturbations were chosen to mimic challenges encountered in practice, such as ambient electrical noise, electrode failures, and sensor displacement in order to test the MU decomposition algorithms sensitivity. Then, the effects of perturbations on the quantity and quality of extracted MUs and neural drive estimation were assessed. Under non-perturbed conditions, an average of 179 ± 40 MUs were extracted. Severe global WGN significantly reduced extracted MUs by approximately 81%. In contrast, more localized WGN, or channel loss as high as 15%, and electrode shift had minimal impact, with reductions in the number of MU decomposed being less than 6%. Reconstruction of neural drive through smoothed cumulative spike trains was significantly impaired by global WGN, thus leading to increased root mean square error when compared to conditions, while localized perturbations had negligible effects. Therefore, the BSS-based MU decomposition methods seem to be robust against localized noise, channel loss, and minor electrode shifts but are vulnerable to global additive noise. These results highlight the importance of carefully applying MU decomposition approaches in practical settings, maintaining high SNRs in EMG recordings, and preprocessing noise treatment to specific MU-decomposition needs.

## 1 Introduction

Surface electromyography (sEMG) signals represent the summation of electrical activity from numerous active motor units in a muscle (Farina et al., 2004; De Luca and Erim, 1994). Each motor unit’s action potentials contribute to the composite EMG, such that the signal contains both a neural component (the motor neuron discharge times) and a peripheral component (the motor unit action potential waveforms) (Negro et al., 2016; Holobar and Zazula, 2007), as shown in Figure 2. Motor unit decomposition (MUD) is a signal processing technique that separates these components and enables the extraction of the firing times of individual motor neurons from the surface EMG (Farina et al., 2004, 2014, 2025). This non-invasive approach provides a direct window into the neural drive to muscles (De Luca and Erim, 1994; Farina and Negro, 2015) and has become a powerful tool for studying motor control and allows researchers to examine motor unit recruitment and firing behavior with high resolution (Del Vecchio et al., 2019a,b; Taleshi et al., 2025a). MUD is essential for applications requiring detailed neural assessments, such as clinical neurophysiology (Gogeascoechea et al., 2020), movement science (Tanzarella et al., 2021), impairment detection and assessment (Valli et al., 2025), rehabilitation (Farina et al., 2017), and studying aging effects (Castronovo et al., 2018). Additionally, surface EMG decomposition serves as a foundational step in developing and training algorithms for real-time applications, providing the necessary data to calibrate and validate online MU decomposition systems (Yeung et al., 2024; Chen et al., 2020; Taleshi et al., 2022; Barsakcioglu and Farina, 2018).

Recent advances in high-density EMG (HD-EMG) electrode grids and decomposition algorithms have improved the reliability and yield of surface MUD by enabling the identification of dozens of motor units from a single contraction (Grison et al., 2025; Nawab et al., 2010). Such high-yield, high-accuracy decomposition has been validated in experimental and simulated studies under ideal conditions, confirming that non-invasive MUD can achieve performance comparable to traditional intramuscular recordings in these scenarios (Nawab et al., 2010).

Despite significant advancements in MU decomposition algorithms, most studies have focused on controlled conditions (de Oliveira et al., 2022; Chen et al., 2020; Lulic-Kuryllo and Inglis, 2022). In typical laboratory experiments, electrode grids are optimally placed and firmly secured, and extensive preprocessing is applied to minimize interference and optimized signal quality. Under these favorable circumstances, decomposition algorithms exhibit robust performance. In real-world settings, however, EMG signals are often subject to significant noise and spatial perturbations that can challenge any algorithm. Namely, changes in skin-electrode impedance result from variations in skin properties due to perspiration, or prolonged electrode wear, causing signal attenuation or distortion (Stegeman et al., 2012; Chen et al., 2025; Clarke et al., 2020). During physical activities, increased sweating alters the conductivity between the skin and electrodes and affects the signal-to-noise ratio. In addition, electrode displacement can occur from natural movements of the skin and underlying muscles during dynamic activities, affecting the spatial consistency of the recorded signals (Muceli et al., 2013). When participants perform tasks involving wrist movements or forearm rotations, the skin shifts relative to the electrodes and leading to misalignment and altered signal acquisition (Muceli et al., 2013). Additionally, poor skin contact, electrode failure, disconnection, or electrode lift in HD-EMG can lead to channel loss (Muceli et al., 2013).

These forms of noise and distortion can disrupt the spatiotemporal patterns that decomposition algorithms rely on, and potentially leads to missed or erroneously identified motor unit discharges. Indeed, it has been shown that increasing the noise level in EMG signals causes a marked decline in the number of motor units that can be accurately identified by current algorithms (Clarke et al., 2020; Chen et al., 2025). Nevertheless, the robustness of contemporary MUD methods to various perturbations remains insufficiently quantified. Many non-invasive motor unit studies have intentionally constrained their analyses to select muscles, subjects, and tasks with high-quality signals, and decomposition algorithms are often tuned or validated on data with minimal noise or interference (de Oliveira et al., 2022). As a result, little is known about how performance deteriorates under non-ideal in vivo conditions compared to controlled experimental settings in which these methods are typically tested. Understanding the sensitivity of MUD to such perturbations is crucial for translating laboratory findings into real-world applications (Farina et al., 2023).

This study quantifies the sensitivity of convolutive blind-source-separation (BSS) based MUD method (Negro et al., 2016) to noises and disturbances in HD-EMG recordings. We introduce eleven perturbation conditions derived from three common noise sources into EMG datasets: (i) additive white Gaussian noise was superimposed on the signals to simulate various levels of background noise, (ii) subsets of EMG channels were set to zero to mimic channel loss or electrode failures, and (iii) spatial shifts of the electrode grid were modeled to mimic electrode displacement during recordings. By applying these perturbations, we evaluate their effects on the quantity and quality of identified motor unit spike trains (MUSTs). In this way, the study provides a systematic assessment of MUD robustness in the face of noise and spatial distortions. Understanding these impacts provides insights into the robustness of current decomposition algorithms under realistic conditions and highlight areas where algorithmic improvements are necessary.

## 2 Methods

### 2.1 Participants

Thirteen healthy adults (four females, nine males; mean age 33 years, range 29–37 years) with no known neurological or musculoskeletal disorders participated in this study. Twelve participants were right-handed, and one was left-handed. Ethical approval was obtained from the Aalto University Ethics Committee, and all participants provided written informed consent prior to the experiment.

### 2.2 Signal Acquisition and Experimental Setup

Simultaneous recordings of joint kinematics and HD-EMG signals were collected from each participant’s dominant arm (Figure 1). HD-EMG signals were acquired using three 8 × 8 electrode arrays (ELSCH064NM3, OT Bioelettronica, Italy), which resulted in a total of 192 channels. Each electrode had a diameter of 4mm with an inter-electrode distance of 10mm. The arrays were positioned to cover the upper half of the forearm, which allowed for the monitoring of most muscles involved in all degrees of freedom (DoFs) of the wrist. Ground and reference electrodes were placed on the wrist.

**Figure 1.**
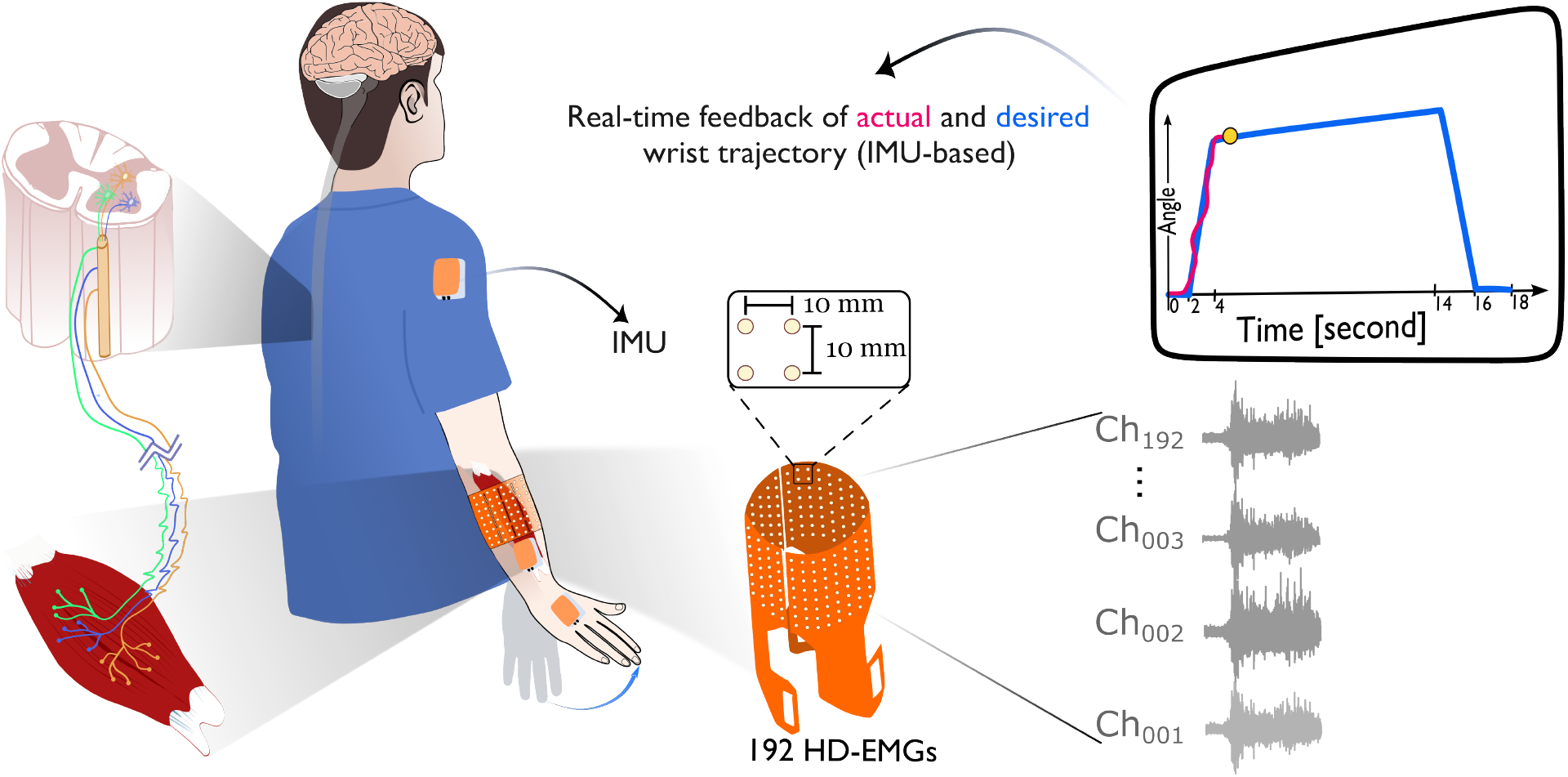
High-density surface EMG setup: three 8 × 8 electrode grids (192 channels total, 4 mm diameter, and 10 mm spacing) cover the upper forearm, with reference and ground electrodes at the wrist. Wireless IMUs on the upper arm, forearm and hand capture joint kinematics simultaneously. Subjects receive realtime visual feedback of their actual versus desired wrist trajectory on a monitor.

The EMG signals were sampled at 2048Hz with 16-bit resolution using a Quattrocento bio-amplifier (OT Bioelettronica, Italy). In-hardware band-pass filtering was applied using a third-order Butterworth filter with cut-off frequencies at 3Hz and 900Hz to remove motion artifacts and high-frequency noise. Joint kinematics were recorded using three wireless inertial measurement units (IMUs) (MTw Awinda, Xsens Technologies B.V., Netherlands) attached to the upper arm, lower forearm, and hand (Figure 1). Measurements of limb movements were obtained from IMUs that were synchronized with the HD-EMG system and sampled at 80Hz.

To ensure high signal quality and maximize the signal-to-noise ratio (SNR), data collection was performed in a magnetically shielded room using a dedicated electrical grid to minimize electromagnetic interference and power supply noise. Participants’ forearms were shaved if necessary and cleaned with alcohol to improve electrode-skin contact. Despite these preparations, due to variations in forearm size or sub-optimal electrode contact in some participants, an average of 15 out of 192 channels (range: 0–42 channels) were excluded from the raw data after visual inspection.

Participants performed three repetitions of each of three wrist motor tasks: flexion/extension, radial/ulnar deviation, and pronation/supination of the forearm. These movements were chosen for their relevance to daily activities and their involvement of multiple wrist DoFs (Youm et al., 1979; Stango et al., 2014). Each task was visually cued using a trapezoidal contraction profile displayed on a screen. The cue included a two-second rest period, a two-second ramp-up phase, and a ten-second steady contraction at a comfortable level that covers the full range of the respective DoF. Real-time feedback of the desired and actual movement trajectories was provided to participants to facilitate accurate task execution Figure 1.

### 2.3 Motor Unit Decomposition (MUD)

During the execution of motor tasks, the electrical activity of activated muscle fibers can be recorded as EMGs. These electrical activities are generated due to the contraction of muscle fibers, and the contraction pattern is regulated by alpha motor neurons (MN) of the spinal cord (De Luca, 1985). This process makes EMG signals a function of the total neural input to the muscle (as shown in the right side of the Figure 1) (Farina et al., 2004). As a result, decomposing EMG signals into their neural components enables the extraction of detailed information about the movement.

In general, multi-channel EMG signals, regardless of the physiological mechanisms, can be explained as a convolution between the discharge timings of motor units (a series of delta functions) and the action potentials of muscle units (with finite duration). This convolutive mixing process underlying EMG generation is shown in Figure 2. The goal in the decomposition algorithms is to reconstruct MUSTs while we have only access to EMGs.

**Figure 2.**
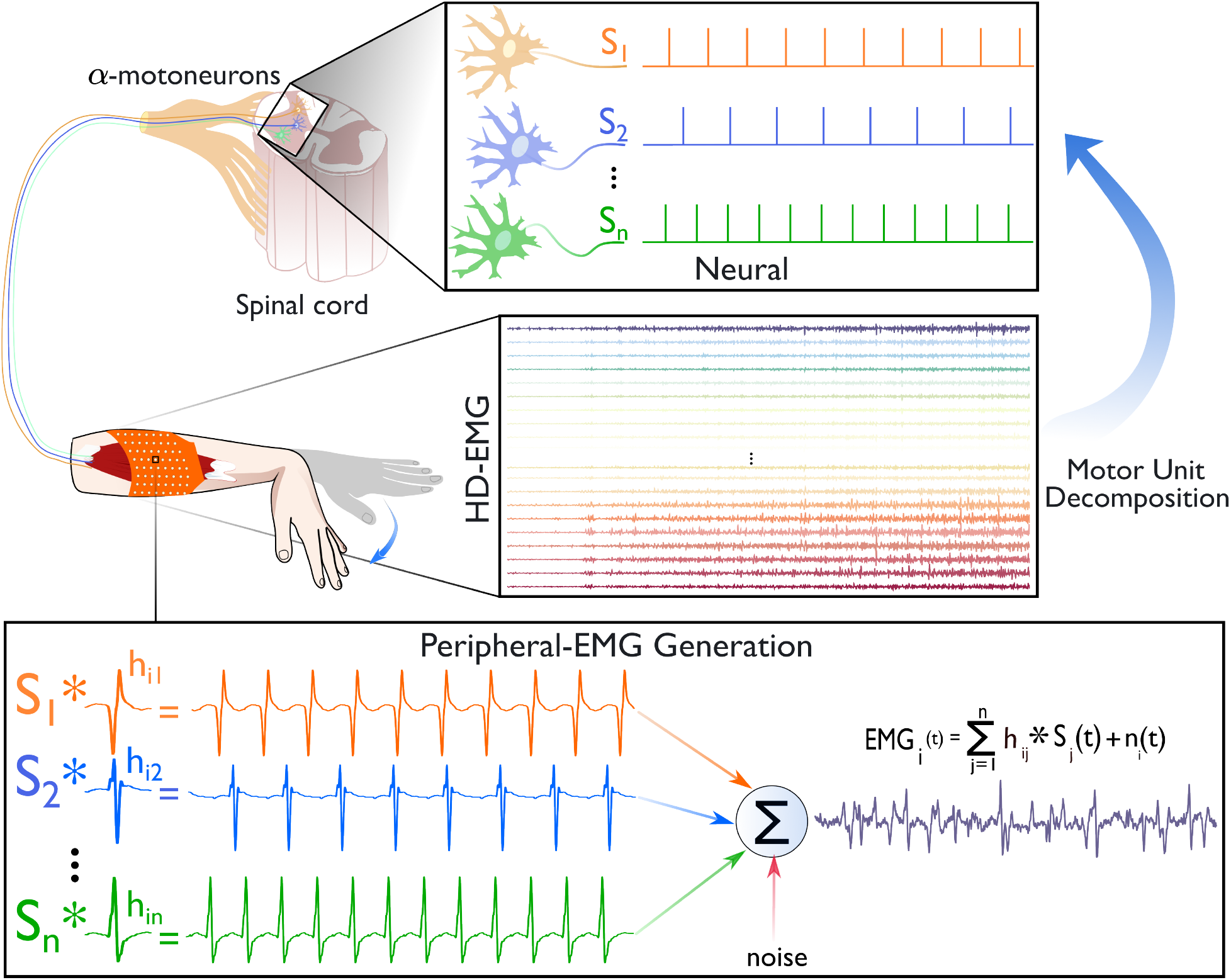
Illustration of the convolutive EMG generation process. A set of n motor unit spike trains *S*_1_ … *S*_*n*_ is convolved with its corresponding motor unit action potential *h*_*ij*_. The measured EMG is the sum of these convolutive mixtures plus noise. Decomposition algorithms aim to recover each *S*_*j*_ from the composite EMG and reveals the precise timing of motor neuron firings. The “*” denotes convolution operation. For simplicity, only a single EMG channel is shown in the lower box of the figure; in practice, motor unit decomposition is applied across all recorded channels.

In this study, MUD was performed using a convolutive BSS algorithm as described in (Negro et al., 2016). To enhance source separation, the multi-channel EMG signals were spatially extended by a factor of E=10 (Taleshi et al., 2022; Yeung et al., 2023a). This extension increases the dimensionality of the data, which improves the identifiability of independent sources in convolutive mixtures. The extended EMG signals were then whitened using singular value decomposition (SVD) (Holobar and Zazula, 2007). This process transforms the signals into a set of uncorrelated components with unit variance. Whitening is a crucial preprocessing step in BSS algorithms, as it simplifies the separation process by reducing signal redundancy and correlational structure. Source estimation was carried out by maximizing the non-Gaussianity and sparsity of the signals. Fixed-point iteration algorithms were employed, incorporating Gram-Schmidt orthogonalization to ensure the uniqueness of the solution (Negro et al., 2016). This approach leverages the inherent statistical properties of motor unit action potentials, which are typically sparse and exhibit nonGaussian distributions. MUSTs were extracted from the squared estimated sources using peak detection and K-means clustering. This process yielded discrete representations of motor unit discharge patterns (Negro et al., 2016). The squaring operation emphasizes the spikes and enhances their detectability against the background noise.

To assess the quality of the decomposition, we employed the silhouette (SIL) measure and the coefficient of variation (CoV) of the inter-spike intervals (ISI). Only motor units with *SIL >* 0.85 (Taleshi et al., 2022, 2025a) and *CoV*_*ISI*_ *<* 30% were accepted for further analysis. The SIL measure provides a normalized indicator of clustering quality. It assesses how well each spike fits within its assigned cluster compared to other clusters. The CoV of ISI evaluates the variability in the firing rates of motor units and serves as a criterion for the physiological plausibility of the discharge patterns. A final manual inspection of the extracted motor units was conducted to verify the reliability of the discharge patterns, as recommended in previous validation studies (Holobar and Zazula, 2007; Negro et al., 2016). This step is essential to confirm the accuracy of the automatic decomposition and to exclude any false units or artifacts from subsequent analyses.

### 2.4 Perturbation Conditions

To assess the sensitivity of motor unit decomposition under simulated real-world challenges, we introduced eleven types of perturbations into the raw EMG signals, derived from three common sources: additive White Gaussian Noise (WGN), channel loss, and electrode shift. These perturbations mimic conditions commonly encountered in practical EMG data collection and aim to evaluate decomposition performance.

#### 2.4.1 White Gaussian Noise (WGN)

Signal noise, particularly additive WGN, can significantly degrade HD-EMG signal quality (Hahne et al., 2017). Factors such as variations in electrode-skin impedance due to drying of the conductive gel (Stegeman et al., 2012; Taleshi et al., 2025b) or perspiration affecting electrode contact (Clarke et al., 2020), can reduce the SNR over time. To simulate these effects, we introduced three WGN perturbation scenarios:

1. *Constant WGN added to all EMG channels*: This simulates a uniform noise level affecting the entire recording and represents ambient electrical noise or consistent degradation of electrode contact.
2. *Progressively increasing WGN on all EMG channels*: Noise level increases over time, modeling scenarios where electrode-skin impedance worsens during prolonged activity due to factors like perspiration.
3. *Constant WGN added to a localized* 4×4 *cluster of channels*: Noise is applied to a randomly selected group of 16 electrodes (a 4×4 cluster) to mimic localized issues such as sweating or degraded electrodeskin contact affecting specific regions of the array.

Each perturbation was applied at three SNR levels: 5dB, 10dB, and 15dB to cover a range from severe to moderate noise conditions. WGN is considered a signal-based perturbation as it alters the raw signal characteristics without changing sensor placement or causing data loss.

#### 2.4.2 Channel Loss

Missing or unusable channels in HD-EMG recordings can occur due to poor electrode-skin contact, disconnections, or low-amplitude signals falling below detection thresholds (Ding et al., 2015; Taleshi et al., 2025b). Such channel loss leads to missing data, which can significantly impact the performance of signal processing and decomposition algorithms. To simulate this condition, we randomly selected approximately 15% of the electrodes (29 out of 192) and set their signals to zero. This reflects real-world scenarios where some channels become nonfunctional, which might affect the extraction of neural information.

#### 2.4.3 Electrode Shift

Electrode displacement is a common issue in surface EMG recordings and often results from movements of the skin relative to the electrode arrays during dynamic activities (Muceli et al., 2013; Taleshi et al., 2025b). This displacement can alter the spatial mapping of muscle activity and degrade the performance of decomposition algorithms. To simulate electrode shift, we artificially shifted the signals from each 8×8 electrode array by one column. This operation makes an eight-channel (1cm) lateral displacement based on the inter-electrode distance.

### 2.5 Evaluation of MUD Under Perturbation Conditions

For each participant, three repetitions of each motion were run through the MUs decomposition algorithm, and clean MUs were extracted. Then, the same MUs decomposition procedure was applied to the EMG signals contaminated with the introduced perturbations described in subsection 2.4.

Rate-of-Agreement (RoA) was used to determine whether motor unit discharge patterns extracted under clean and perturbed conditions corresponded to the same motor units Negro et al. (2016). For the *j*^th^ motorunit source, RoA was defined as RoA_*j*_ = (*Q*_Cln∩Prt, *j*_*/Q*_Cln∪Prt, *j*_) × 100% where *Q*_Cln∩Prt, *j*_ is the number of discharge events detected in both conditions (with a timing tolerance of ± 0.5 ms) and *Q*_Cln∪Prt, *j*_ is the total number of unique discharge events detected in either condition. Following Holobar and Zazula (2007), two spike trains were considered to originate from the same motor unit when the common discharges accounted for more than 30% of the total discharges in each train. This threshold ensures sufficient overlap between spike trains to confidently associate them with the same motor unit.

In addition, to evaluate the effect of noise on the estimation of neural drive to muscles, we analyzed the smoothed cumulative spike trains (sCSTs) derived from MUSTs under both clean and perturbed conditions. The neural drive was estimated by summing binary spike trains across all motor units to obtain the cumulative spike train (CST). The CSTs were smoothed to obtain the sCSTs, with the goal of reducing variability and emphasising the underlying neural drive patterns.

Smoothing was performed by applying a moving average with a window length of 100ms and a 50% overlap, corresponding to 205 samples at the EMG sampling rate of 2048Hz. To further reduce high-frequency fluctuations, we applied a Gaussian filter using MATLAB’s “smoothdata” function with a window size of 20 samples and the “gaussian” method. The smoothed CSTs were then interpolated back to the original time scale using MATLAB’s “interp1” function with the “spline” method to ensure temporal alignment with the original CSTs.

To assess the impact of noise, the sCSTs obtained from the clean EMG data were compared with those from the perturbed EMG data under various noise conditions, as described in subsection 2.4. The similarity between the sCSTs from clean and noisy conditions was quantified using the root mean square error (RMSE). This metric provided insights into how different types of noise affect the neural drive estimation.

### 2.6 Statistical Analysis

Statistical analyses were conducted to assess significant differences in MUD under perturbation conditions. The Kolmogorov-Smirnov test was employed to verify the normality of the data distributions. Paired ttests were used to compare the number of extracted motor units between clean and perturbed conditions. We reported the mean and 95% confidence intervals (CIs) of the number of extracted motor units. The confidence interval provides an estimated range that is likely to contain the true population mean. We also calculated the percentage change in the number of extracted motor units between clean and perturbed conditions. Statistical significance was determined at a threshold of p<0.05. All analyses were performed using Python’s SciPy library.

## 3. Results

### 3.1 Impact of Noise on MU Extraction and Matching

Table 1 shows the average number of accurately extracted MUs across different conditions, along with the number of matched MUs by RoA analysis. In the clean condition, an average of 179±40 MUs were extracted across all motions, with 2637 MUs per motion. Notably, extension and ulnar deviation showed higher extraction numbers than other motions.

**Table 1.**
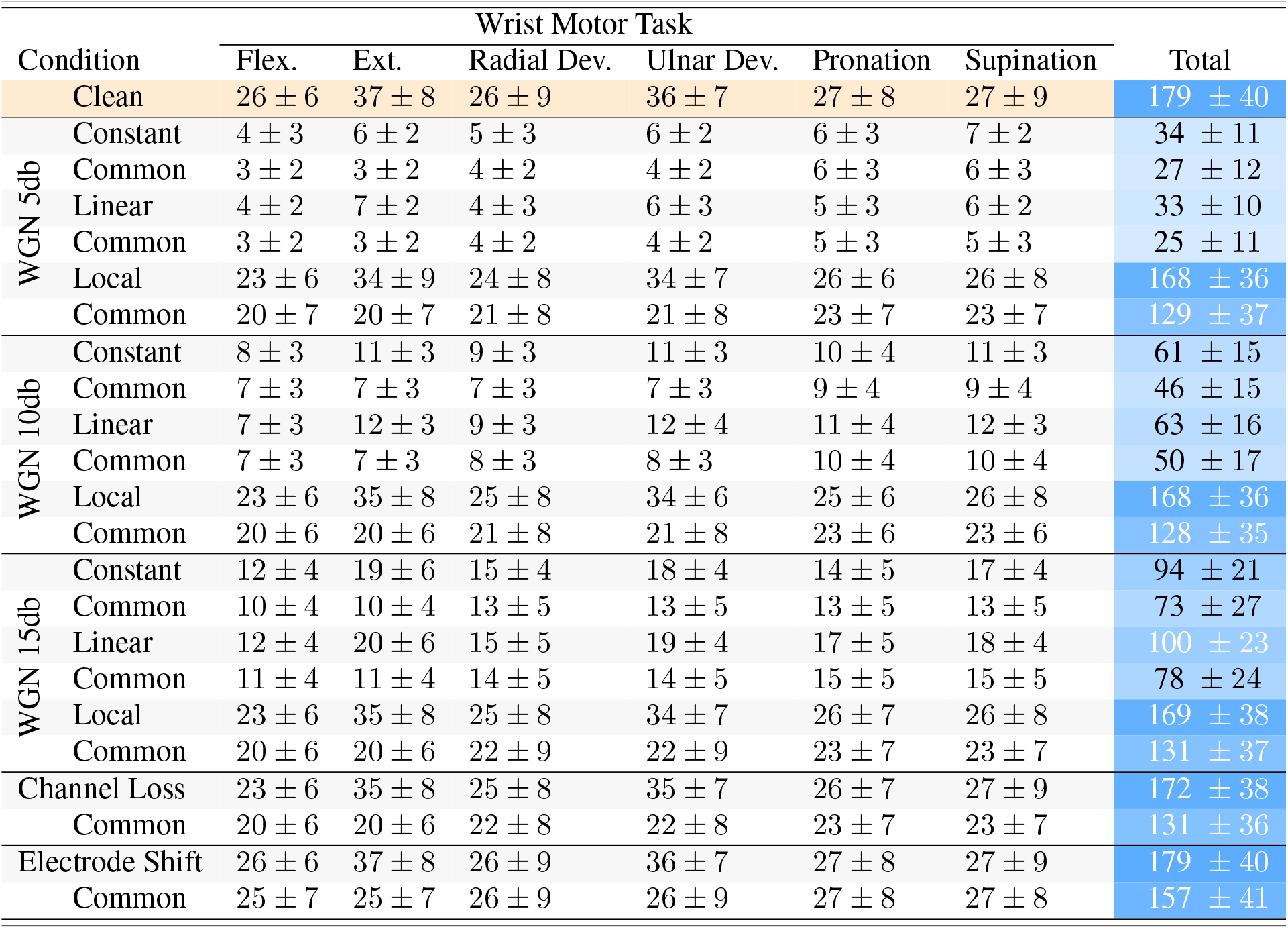
Motor unit decomposition performance for 13 subjects under clean and 11 degraded conditions. The number of identified motor units and match counts are reported as Mean ± CI.

The addition of constant and linear WGN at 5 dB caused the most severe degradation. The total number of accurately identified units fell to 34±11 (81 %) and 33±10 (82 %) MUs relative to the clean condition, respectively. The RoA matched 27±12 and 25±11 of these units, which correspond to matching rates of 79 % and 76 %. MUD maintained a high extraction performance under Local WGN at the same SNR with 168±36 units were identified, only 6 % below clean condition. The number of matched MUs was 129±37 (77 %).

At 10 dB, constant and linear WGN still caused a significant drop in the identified number of MUs. MUD identified 61±15 and 63±16 MUs (66 % and 65 % reductions). The numbers of matched units rose to 46±15 and 50±17, which equivalent to 7579 % matching rates. Local WGN again produced near-clean performance with identifying 168±36 units and 128±35 (76 %) matches.

An SNR of 15 dB further mitigated the effect of constant and linear WGN. The MUD algorithm estimated 94±21 and 100±23 MUs units (reductions of 47 % and 44 %). The number of matched MUs increased to 73±27 and 78±24, with matching rates around 78%. Local WGN remained almost neutral as the MUD identified 169±38 MUs with 131±37 matches (77 %).

15% channel loss resulted in a total of 172±38 MUs extracted, only 4 % below clean condition. The number of matched MUs was 131±36, with a matching rate of about 76%. Similarly, the simulated 1 cm electrode shift showed a negligible effect on MU extraction. The total number of extracted MUs remained consistent with the clean condition at 179±40 and raised the matching rate to 88 % (157±41). A paired *t*-test confirmed that constant and linear WGN at every SNR significantly decreased the number of extracted MUs (*p <* 0.05),

### 3.2 Noises Impact on Neural Derive

To assess the effect of noise on neural drive estimation, the similarity between sCSTs under clean and noisy conditions was quantified using RMSE as shown in Figure 3. The bar plots indicate the mean RMS error values with 95% confidence intervals (CI), while pink points show individual participant data. For constant WGN at 5 dB, the RMSE ranged from 9.25 (Radial Deviation) to 19.58 (Extension), with an average RMSE of 10.69 across motions. The linear WGN at the same dB level produced slightly lower RMSE values, with an average of 10.39 RMSE. In contrast, localized WGN at 5 dB showed significantly lower RMSE values across all motions, with an average RMSE of 1.43.

**Figure 3.**
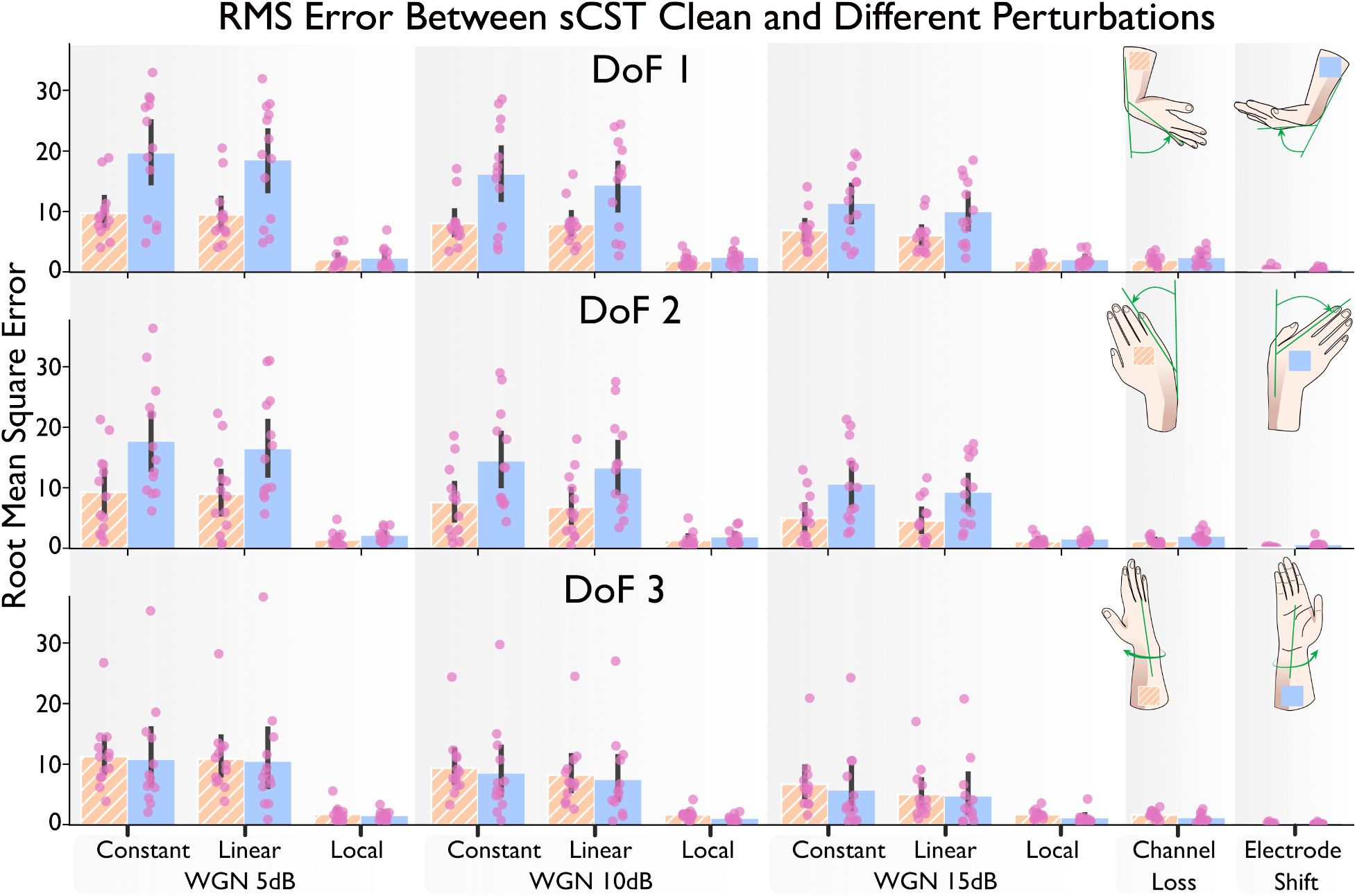
Root mean square (RMS) error between smoothed cumulative spike trains (sCST) under clean and 11 perturbed conditions across degrees of freedom (DoF). The bar plots shows the RMS error for sCST estimation under 11 noise and perturbation conditions. The noise conditions are Constant, Linear, and Local White Gaussian Noise (WGN) at 5 dB, 10 dB, and 15 dB, as well as Channel Loss and Electrode Shift. Each row shows distinct movement directions and corresponds to a different DoF: DoF 1, DoF 2, and DoF 3. Orange hatched bars denote the positive movement direction (+DoF, including Flexion, Radial Deviation, and Pronation), and blue bars denote the opposite direction (-DoF, including Extension, Ulnar Deviation, and Supination), as illustrated by the hand icons. Bars plots indicate mean RMS error values with 95% confidence interval (CI). Pink points show individual participants.

As the noise level increased to 10 dB, the RMSE under constant WGN showed a marked decrease across all motions, with an average RMSE of 8.43, while linear WGN resulted in an average of 7.37 RMSE, and localized WGN condition showed an average 0.97 RMSE. At 15 dB, the constant WGN led to an average RMSE of 5.61, reflecting an approximate reduction of 47.46% in accuracy compared to 5 dB. The linear WGN condition showed a similar trend, with a 55.36% decrease (from 10.39 to 4.64 RMSE) in accuracy. However, the localized WGN showed an average RMSE of 1.03, consistent with previous noise levels. A15% channel loss and 1 cm electrode shift showed even lower RMSE values of 1.08 and 0.13%, respectively.

Across individual motions, the noise impact varied, with the highest effects observed in Extension and Ulnar Deviation, which showed average RMSEs of 8.95 and 8.08, respectively. Flexion and Pronation followed, with RMSEs of 5.07 and 5.29, showing moderate sensitivity to perturbations. Finally, Radial Deviation and Supination were the least affected, with RMSE values of 4.29 and 4.71, respectively.

## 4 Discussion

This study examined the sensitivity of convolutive BSS based MU decomposition methods to common perturbations. By introducing WGN, channel loss, and electrode shift, we assessed their impact on the quantity and quality of extracted MUSTs and the estimation of neural drive to muscles. The present findings show that (i) global additive noise (uniform WGN) is the single most disruptive perturbation; once the overall SNR approaches 5–10 dB the number of identifiable motor units and the fidelity of the neural-drive estimate decline sharply (ii) local disturbances, whether they are restricted noise patches, the loss of ∼15% of channels, or a 1 cm grid shift, have only a marginal effect on both yield and neural-drive metrics (iii) the impact of a given perturbation is task-dependent: extension and ulnar deviation are slightly more sensitive than the other four wrist movements.

### 4.1 Motor Units Decomposition Sensitivity to Perturbations

MUD identified an average of 179±40 MUs for all motions during clean condition. Our results showed that moderate levels of additive WGN had only a limited effect on decomposition yield and quality, whereas severe noise caused a pronounced effect. Specifically, during constant and linear WGN at SNR levels of 5 dB and 10 dB, the number of accurately extracted MUs significantly dropped to 81-82% and 66-65%, respectively. At an SNR of 15 dB, the impact of constant and linear WGN was less severe, with reductions of approximately 44-47% in extracted MUs. This behavior is consistent with the inherent noise tolerance of convolutive BSS algorithms (Yeung et al., 2024). For instance, the convolution kernel compensation (CKC) approach has been reported to tolerate high levels of additive noise while still detecting individual motor unit firings (Zazula et al., 2006). In line with this, we observed that signal quality began to seriously impair motor unit identification only when the global SNR dropped to around 5 dB. At this extreme SNR, the algorithm’s performance deteriorated significantly, which is an expected outcome, since the motor unit action potentials (MUAPs) are barely distinguishable from the noise floor in such conditions. Indeed, recent analyses have confirmed that WGN is the dominant noise factor influencing sEMG decomposition performance, far more than other noise sources like baseline wander or powerline interference (Zhao et al., 2024). Thus, our finding of significant degradation at 5 dB and 10 db SNR is in agreement with the broader literature when the overall noise power approaches the level of MUAP signals, the blind source separation process can no longer reliably isolate the underlying sources. These results highlight the critical importance of maintaining high SNR during EMG recordings to ensure the reliability of MU decomposition. In practical terms, this emphasizes the need for meticulous electrode preparation, such as skin cleaning and abrasion, use of conductive gels, and minimization of external electrical interference, to maximize signal quality.

In contrast to global WGN noise, the BSS-based decomposition proved relatively robust to localized disturbances such as noise bursts on a 4×4 channels or the loss of a subset (15%) of channels. Even at localized 5 dB, the number of extracted MUs falls only by 6%. Similarly, with 15% of channel loss, the MUD can identify 172±38 units, 76% of which match the clean set. The possible technical reason for this resilience lies in the high spatial sampling of HD-sEMG (Drost et al., 2006). Each motor unit’s action potential is detected by many neighboring electrodes, so information redundancy is built into the recordings (Drost et al., 2006). Even if one channel is heavily noisy or fails, its neighbors still capture the MUAP waveform and allows the source separation to proceed with minimal impact. Our results reflect this mechanism where the MUD performance dropped only marginally when a cluster of channels was compromised and confirms that the convolutive BSS approach can leverage the spatial redundancy to fill in missing or noisy data. This robustness is encouraging for real-world use, as it implies that sporadic electrode failures or interference on a few contacts will not severely degrade the overall decoding of the neural signals.

A one-column (1 cm) lateral or medial displacement of the HD-sEMG grid had virtually no impact on decomposition performance. Specifically, the convolutive BSS algorithm extracted 179±40 motor units and 88% of those units matched the clean condition. This invariance can be attributed to two complementary factors. First, the algorithm operates on spatially extended EMG observations and applies whitening and fixed-point source extraction that are insensitive to electrode ordering (Negro et al., 2016). Shifting the entire column changes channel indices but not the underlying spatial information, so the whitening matrix and subsequent separation remain valid (Negro et al., 2016). Second, with a 10 mm inter-electrode distance the MUAP generated by a given motor unit is still sampled by several neighbouring electrodes after a 1 cm shift. As a result, the convolutional mixing filters remain well represented in the signal (Del Vecchio et al., 2020). Together, these properties preserve both the sparsity and the spatial independence exploited by BSS, preventing any appreciable loss of motor-unit yield or neural-drive fidelity.

Regarding the impact on neural drive estimation, global WGN significantly increased the RMS error between the smoothed CST from clean and noisy conditions. Under 5 dB constant WGN, neural drive estimation is substantially impaired, as evidenced by an average RMSE of 10.69. As the signal became cleaner (with increasing SNR), the RMSE values fell, which suggests enhanced preservation of the neural drive signal. Moreover, localized WGN at the same SNR levels had minimal effects on the RMSE, with average values as low as 1.43 at 5 dB, close to the clean condition. A 15% channel loss and 1 cm electrode shift resulted in RMSE values of 1.08 and 0.13, respectively. These low values suggest a negligible impact of these noises on neural drive estimation.

We also observed some differences in decomposition outcomes across various wrist motion tasks. Certain dynamic or complex movements resulted in a slightly lower yield of identifiable motor units compared to simpler or more static tasks. Notably, during wrist extension and ulnar deviation, a higher number of MUs were identified. This task-dependent variations might be due to activation of different subsets and patterns of motor units for each movement (Warriner et al., 2022). At the same time, wrist extension and ulnar deviation might be involve more distinctly isolated surface muscles and this makes MU identification easier.

Alternative explanations for the observed robustness to localized perturbations could involve the inherent properties of the BSS algorithm and the spatial distribution of MUs. Since MU action potentials are spatially distributed across the electrode array, localized noise or channel loss may not significantly disrupt the overall decomposition if sufficient spatial information is preserved (Disselhorst-Klug et al., 1999). However, global noise affects all channels uniformly, severely degrading the spatial and temporal cues necessary for separation. The redundancy in HD-sEMG systems allows for compensation when only a subset of channels is affected, but when all channels are contaminated, the algorithm’s ability to isolate individual MUs is compromised.

### 4.2 Limitations and Future work

While this study provides a thorough analysis of noise and perturbation sensitivity, several limitations must be acknowledged. First, our evaluation of decomposition performance under noise was primarily quantitative. The ground truth for motor unit firings in real sEMG data is generally unknown, so we relied on performance metrics. There is an implicit assumption that if the algorithm outputs fewer motor units at 5 dB SNR, it indeed reflects lost motor unit information. In future work, it would be valuable to include complementary validation techniques, such as cross-comparing the decomposition output with simultaneously recorded intramuscular EMG spike trains or using multiple decomposition algorithms and checking for consensus (Yeung et al., 2023b, 2024). Second, although we identified that the algorithm is resilient to many perturbations, we did not explicitly test combinations of adverse factors. In practice, multiple challenges may occur simultaneously. Designing experiments to systematically combine factors would help map out the boundaries of the algorithm’s robustness. Third, the perturbations were artificially introduced and may not capture the full complexity of noise encountered in real-world conditions, such as motion artifacts, electromagnetic interference, or physiological variations among individuals.

Future research should focus on enhancing the resilience of MU decomposition algorithms to global noise. The enhanced robustness could be achieved by investigating adaptive decomposition methods (Chen et al., 2020; Yeung et al., 2024). This approach may lead to more precise and stable MUs. In addition, higher robustness might be achieved through adaptive filtering, machine learning approaches, or advanced signal processing techniques that can differentiate between noise and signal components. Furthermore, this could involve improved electrode designs, better skin preparation techniques, or the incorporation of real-time noise reduction methods.

### 4.3 Conclusion

Our findings show that convolutive BSS-based MU decomposition is resilient to localized noise, moderate channel loss, and small electrode shifts, yet highly sensitive to global additive noise. Across 13 participants and six wrist degrees of freedom, localized perturbations or 15% channel loss reduced the number of accurately identified motor units by less than 6%, and neural-drive estimates stayed nearly unchanged. In contrast, global WGN at 5 dB SNR cut the yield by about 81% and markedly degraded cumulative spike-train fidelity. These results underscore the need to maintain high signal-to-noise ratios during HD-sEMG acquisition and suggest that future algorithmic and hardware refinements should prioritize robustness against pervasive broadband noise to enable reliable, real-world motor-unit decomposition.

## Conflict of Interest

The authors declare that there are no conflicts of interest, financial or otherwise, associated with this manuscript.

### Acknowledgment

The authors are grateful to the Aalto Science-IT project for the provision of computational resources. The cluster computing tools supplied by this project were instrumental in facilitating the decomposition and analysis of motor units for our study.

## Data Availability Statement

Data are available from the corresponding authors upon reasonable request.

## Author Contributions

All authors have discussed the results of the study and approved the final version of the manuscript. Mansour Taleshi: conceptualization, methodology, data analysis, visualization, writing original and final draft; Dennis Yeung: experiments; Francesco Negro: final draft; Ivan Vujaklija: conceptualization, methodology, revision and editing, final draft.

## References

Barsakcioglu, D. Y. and Farina, D. (2018). A real-time surface emg decomposition system for non-invasive human-machine interfaces. In 2018 IEEE Biomedical Circuits and Systems Conference (BioCAS), pages 1–4. IEEE.

Castronovo, A. M., Mrachacz-Kersting, N., Stevenson, A. J. T., Holobar, A., Enoka, R. M., and Farina, D. (2018). Decrease in force steadiness with aging is associated with increased power of the common but not independent input to motor neurons. Journal of neurophysiology, 120(4):1616–1624.

Chen, C., Li, D., and Xia, M. (2025). A motor unit action potential-based method for surface electromyography decomposition. Journal of NeuroEngineering and Rehabilitation, 22(1):1–16.

Chen, C., Ma, S., Sheng, X., Farina, D., and Zhu, X. (2020). Adaptive real-time identification of motor unit discharges from non-stationary high-density surface electromyographic signals. IEEE Transactions on Biomedical Engineering, 67(12):3501–3509.

Clarke, A. K., Atashzar, S. F., Del Vecchio, A., Barsakcioglu, D., Muceli, S., Bentley, P., Urh, F., Holobar, A., and Farina, D. (2020). Deep learning for robust decomposition of high-density surface emg signals. IEEE Transactions on Biomedical Engineering, 68(2):526–534.

De Luca, C. J. (1985). Control properties of motor units. Journal of Experimental Biology, 115(1):125–136.

De Luca, C. J. and Erim, Z. (1994). Common drive of motor units in regulation of muscle force. Trends in neurosciences, 17(7):299–305.

de Oliveira, D. S., Casolo, A., Balshaw, T. G., Maeo, S., Lanza, M. B., Martin, N. R., Maffulli, N., Kinfe, T. M., Eskofier, B. M., Folland, J. P., et al. (2022). Neural decoding from surface high-density emg signals: influence of anatomy and synchronization on the number of identified motor units. Journal of Neural Engineering, 19(4):046029.

Del Vecchio, A., Casolo, A., Negro, F., Scorcelletti, M., Bazzucchi, I., Enoka, R., Felici, F., and Farina, D. (2019a). The increase in muscle force after 4 weeks of strength training is mediated by adaptations in motor unit recruitment and rate coding. The Journal of physiology, 597(7):1873–1887.

Del Vecchio, A., Holobar, A., Falla, D., Felici, F., Enoka, R., and Farina, D. (2020). Tutorial: Analysis of motor unit discharge characteristics from high-density surface emg signals. Journal of Electromyography and Kinesiology, 53:102426.

Del Vecchio, A., Negro, F., Holobar, A., Casolo, A., Folland, J. P., Felici, F., and Farina, D. (2019b). You are as fast as your motor neurons: speed of recruitment and maximal discharge of motor neurons determine the maximal rate of force development in humans. The Journal of physiology, 597(9):2445–2456.

Ding, Q., Han, J., Zhao, X., and Chen, Y. (2015). Missing-data classification with the extended full-dimensional gaussian mixture model: applications to emg-based motion recognition. IEEE Transactions on Industrial Electronics, 62(8):4994–5005.

Disselhorst-Klug, C., Rau, G., Schmeer, A., and Silny, J. (1999). Non-invasive detection of the single motor unit action potential by averaging the spatial potential distribution triggered on a spatially filtered motor unit action potential. Journal of Electromyography and Kinesiology, 9(1):67–72.

Drost, G., Stegeman, D. F., van Engelen, B. G., and Zwarts, M. J. (2006). Clinical applications of high-density surface emg: a systematic review. Journal of Electromyography and Kinesiology, 16(6):586–602.

Farina, D., Merletti, R., and Enoka, R. M. (2004). The extraction of neural strategies from the surface emg. Journal of applied physiology, 96(4):1486–1495.

Farina, D., Merletti, R., and Enoka, R. M. (2014). The extraction of neural strategies from the surface emg: an update. Journal of applied physiology, 117(11):1215–1230.

Farina, D., Merletti, R., and Enoka, R. M. (2025). The extraction of neural strategies from the surface emg: 2004–2024. Journal of Applied Physiology, 138(1):121–135.

Farina, D. and Negro, F. (2015). Common synaptic input to motor neurons, motor unit synchronization, and force control. Exercise and sport sciences reviews, 43(1):23–33.

Farina, D., Vujaklija, I., Brånemark, R., Bull, A. M., Dietl, H., Graimann, B., Hargrove, L. J., Hoffmann, K.-P., Huang, H., Ingvarsson, T., et al. (2023). Toward higher-performance bionic limbs for wider clinical use. Nature biomedical engineering, 7(4):473–485.

Farina, D., Vujaklija, I., Sartori, M., Kapelner, T., Negro, F., Jiang, N., Bergmeister, K., Andalib, A., Principe, J., and Aszmann, O. C. (2017). Man/machine interface based on the discharge timings of spinal motor neurons after targeted muscle reinnervation. Nature biomedical engineering, 1(2):1–12.

Gogeascoechea, A., Kuck, A., Van Asseldonk, E., Negro, F., Buitenweg, J. R., Yavuz, U. S., and Sartori, M. (2020). Interfacing with alpha motor neurons in spinal cord injury patients receiving trans-spinal electrical stimulation. Frontiers in neurology, 11:493.

Grison, A., Mendez Guerra, I., Clarke, A. K., Muceli, S., Ibáñez, J., and Farina, D. (2025). Unlocking the full potential of high-density surface emg: novel non-invasive high-yield motor unit decomposition. The Journal of Physiology.

Hahne, J. M., Markovic, M., and Farina, D. (2017). User adaptation in myoelectric man-machine interfaces. Scientific reports, 7(1):1–10.

Holobar, A. and Zazula, D. (2007). Multichannel blind source separation using convolution kernel compensation. IEEE Transactions on Signal Processing, 55(9):4487–4496.

Lulic-Kuryllo, T. and Inglis, J. G. (2022). Sex differences in motor unit behaviour: A review. Journal of Electromyography and Kinesiology, 66:102689.

Muceli, S., Jiang, N., and Farina, D. (2013). Extracting signals robust to electrode number and shift for online simultaneous and proportional myoelectric control by factorization algorithms. IEEE Transactions on Neural Systems and Rehabilitation Engineering, 22(3):623–633.

Nawab, S. H., Chang, S.-S., and De Luca, C. J. (2010). High-yield decomposition of surface emg signals. Clinical neurophysiology, 121(10):1602–1615.

Negro, F., Muceli, S., Castronovo, A. M., Holobar, A., and Farina, D. (2016). Multi-channel intramuscular and surface emg decomposition by convolutive blind source separation. Journal of neural engineering, 13(2):026027.

Stango, A., Negro, F., and Farina, D. (2014). Spatial correlation of high density emg signals provides features robust to electrode number and shift in pattern recognition for myocontrol. IEEE Transactions on Neural Systems and Rehabilitation Engineering, 23(2):189–198.

Stegeman, D. F., Kleine, B. U., Lapatki, B. G., and Van Dijk, J. P. (2012). High-density surface emg: Techniques and applications at a motor unit level. Biocybernetics and biomedical engineering, 32(3):3–27.

Taleshi, M., Bubeck, F., Brunner, P., Gizzi, L., and Vujaklija, I. (2025a). Observing changes in motoneuron characteristics following distorted sensorimotor input via blood flow restriction. Journal of Applied Physiology, 138(2):559–570.

Taleshi, M., Yeung, D., Dinh Trong, M., Negro, F., Deny, S., and Vujaklija, I. (2025b). Impact of noise on deep learning-based pseudo-online gesture recognition with high-density emg. bioRxiv.

Taleshi, M., Yeung, D., Negro, F., and Vujaklija, I. (2022). Muscle synergy-driven motor unit clustering for human-machine interfacing. In 2022 44th Annual International Conference of the IEEE Engineering in Medicine & Biology Society (EMBC), pages 4147–4150. IEEE.

Tanzarella, S., Muceli, S., Santello, M., and Farina, D. (2021). Synergistic organization of neural inputs from spinal motor neurons to extrinsic and intrinsic hand muscles. Journal of Neuroscience, 41(32):6878–6891.

Valli, G., Sarto, F., Negro, F., Monti, E., Sirago, G., Paganini, M., Zampieri, S., Franchi, M. V., Casolo, A., Candia, J., et al. (2025). Changes in motor unit conduction velocity after unilateral lower limb suspension and active recovery correlate with muscle ion channel gene expression. bioRxiv, pages 2025–02.

Warriner, C. L., Fageiry, S., Saxena, S., Costa, R. M., and Miri, A. (2022). Motor cortical influence relies on task-specific activity covariation. Cell reports, 40(13).

Yeung, D., Negro, F., and Vujaklija, I. (2023a). Optimal motor unit subset selection for accurate motor intention decoding: Towards dexterous real-time interfacing. IEEE Transactions on Neural Systems and Rehabilitation Engineering.

Yeung, D., Negro, F., and Vujaklija, I. (2023b). Semi-automated identification of motor units concurrently recorded in high-density surface and intramuscular electromyography. In 2023 45th Annual International Conference of the IEEE Engineering in Medicine & Biology Society (EMBC), pages 1–5. IEEE.

Yeung, D., Negro, F., and Vujaklija, I. (2024). Adaptive hd-semg decomposition: towards robust real-time decoding of neural drive. Journal of Neural Engineering, 21(2):026012.

Youm, Y., Dryer, R., Thambyrajah, K., Flatt, A., and Sprague, B. (1979). Biomechanical analyses of forearm pronation-supination and elbow flexion-extension. Journal of Biomechanics, 12(4):245–255.

Zazula, D., Holobar, A., and Divjak, M. (2006). Convolution kernel compensation applied to 1d and 2d blind source separation. In International Conference on Security and Cryptography, volume 2, pages 126–133. SCITEPRESS.

Zhao, Z., Guo, W., Xu, Y., and Sheng, X. (2024). A biosignal quality assessment framework for high-density semg decomposition. Biomedical Signal Processing and Control, 90:105800.

